# Distributed Biomarker Discovery for Immuno-Oncology

**DOI:** 10.1101/2025.07.25.666789

**Authors:** Farnoosh Abbas-Aghababazadeh, Kewei Ni, Minoru Nakano, Xin Wang, Sisira Kadambat Nair, Nasim Bondar Sahebi, Nadir Sella, Sofija Spasojevic, Jonas Denck, Nicolas Riesterer, Thomas Lehéricy, Auranuch Lorsakul, Ramtin Zargari Marandi, Sina Nassiri, Antoaneta Vladimirova, Ruben Armañanzas, John Stagg, Magnus Fontes, Cameron MacPherson, Benjamin Haibe-Kains

## Abstract

**Background:** Identifying robust predictive biomarkers for immuno-oncology (IO) therapy response remains challenging due to the complexity of tumor host interactions and the limited availability of public datasets. While achieving clinical relevance requires biomarkers that generalize across diverse datasets, privacy concerns surrounding sensitive clinical and molecular data further restrict data sharing, impeding progress in biomarker discovery.

**Materials and Methods:** We developed a scalable, cloud-based distributed pipeline, implemented as an *in-silico* simulation of federated learning, to enable privacy-preserving integration of clinical and molecular data across institutions. This approach supports the development of genomic prediction algorithms without sharing sensitive patient data. Using curated gene expression profiles, we evaluated the association of tumor microenvironment and IO gene expression signatures with clinical outcomes across pan-cancer, cancer-specific, and treatment-specific settings, and developed a multivariable distributed model to predict IO response.

**Results:** We implemented an integrated meta-analysis pipeline using harmonized data across 18 datasets comprising 967 patients and seven cancer types treated with PD-1/PD-L1, CTLA-4, or combination immunotherapies. This approach identified biomarkers specific to lung cancer and CTLA-4 therapy previously unreported in pan-cancer and pan-IO analyses, highlighting the value of context-specific associations. Similar gene expression signatures were associated with progression-free survival and response in both pan-cancer and melanoma datasets, with notable overlap between dual checkpoint blockade and PD-1/PD-L1 therapies. Distributed multivariable XGBoost models outperformed the average of locally trained models, indicating improved generalizability.

**Conclusion:** We present a new distributed pipeline for biomarker discovery in immuno-oncology that preserves patient privacy through federated analysis and secure data handling. This study highlights the importance of larger, diverse datasets to refine cancer- and treatment-specific biomarkers, paving the way for more precise and personalized IO therapies.

## Introduction

Immuno-oncology (IO) therapies, such as checkpoint inhibitors targeting programmed cell death protein-1 (PD-1), programmed death-ligand 1 (PD-L1), and cytotoxic T-lymphocyte-associated protein 4 (CTLA-4), have revolutionized cancer treatment by providing long-lasting responses in some patients with advanced cancers. However, a substantial number of patients fail to respond, and even those who initially benefit often develop resistance [1]. Furthermore, immunotherapy is not without risk, with a spectrum of mild to life-threatening immune-related adverse events. Identifying patients unlikely to benefit from IO therapy can also shed light on the resistance mechanisms and assist with clinical decision-making. The effectiveness of these therapies varies by cancer type, making accurate patient selection a critical clinical unmet need.

Investigating the mechanisms of immune resistance has created substantial opportunities to identify biomarkers that predict patient response to IO therapy. While several biomarkers have been validated for clinical use, they remain limited. These include the expression of target ligands such as PD-L1 [2], co-inhibitory receptors such as PD-1 [3] and LAG3 [4], tumor mutational burden (TMB) [≥10 mutations/Mb] [5], tumor-infiltrating lymphocytes (TIL) [6], microsatellite instability (MSI) [7,8], T-cell repertoire [9], and secretion of IFNγ [10]. However, these biomarkers have low concordance and rarely translate to more than a few cancer subtypes. In most cancers, IO therapies are administered without predictive biomarker assessment, resulting in subsets of patients incurring unnecessary toxicity and deriving limited clinical benefit.

The mechanisms by which intercellular signaling pathways modulate sensitivity or resistance to immunotherapy remain inadequately understood, and existing biomarkers fail to capture these biological signals [11]. For instance, PD-L1 expression is hampered by arbitrary cut-off thresholds and assay discordance, particularly in combination therapies [12]. Similarly, TMB lacks standardization due to variability in sequencing technologies and bioinformatics processing, and cancer-specific TMB thresholds may be needed to account for heterogeneity across tumor types [13]. Moreover, given the complex interactions within the tumor microenvironment (TME), developing biomarkers that capture these interactions could potentially improve the ability to identify patients who are most likely to benefit from IO therapies [14,15].

Multiple gene expression (GE) signatures have been proposed as biomarkers for IO therapy [16–18]. However, most are derived from single tumor types and small patient datasets, leading to underpowered analyses that ultimately fail to accurately identify clinically-meaningful and reproducible biomarkers [19,20]. Consequently, few predictive biomarkers of IO response and intrinsic IO resistance mechanisms have been discovered [21–24]. Recently, we and others reported large meta-analyses of previously described biomarkers for checkpoint inhibitor response [25–28]. Robust biomarker discovery requires access to large, diverse datasets, but progress is often limited by the lack of publicly available genomic, transcriptomic, and clinical data from clinical trials.

Access to data is often hindered by patient privacy concerns, high management and storage costs, restrictive sharing policies, and complex agreements with pharmaceutical companies and institutions to ensure compliance with privacy regulations [29–31].

Multi-institutional collaborations present a promising solution, traditionally relying on a centralized data-sharing paradigm. However, sharing sensitive clinical trial data remains challenging due to regulatory hurdles and importance of preserving patient confidentiality. Even anonymization methods are often insufficient to guarantee privacy [32]. Federated Learning (FL) offers an innovative alternative by adopting a model-to-data approach, enabling multi-institutional analyses without requiring data to be pooled in a central repository, thereby addressing data governance and privacy concerns [33]. FL enables each institution to maintain its own data governance policies and control data access. Institutions retain their own computing and data storage facilities, sharing only models across organizations to construct global models. By preserving data control and ownership while fostering collaboration, this approach has the potential to improve and scale biomedical research, which would be highly beneficial for IO biomarker discovery. It would allow scientists to leverage sufficiently large data compendia to develop more effective predictors of IO response, identify patients who will benefit from IO therapy, understand resistance mechanisms, reduce treatment costs, and advance precision medicine in cancer immunotherapy. However, the complexity of deploying FL in real-world institutional networks has limited its progress, confining most FL studies to simulations thus far [34,35].

In this study, we implemented a streamlined version of the FL framework as an *in-silico* simulation, adapting meta-analysis and multivariable prediction within a distributed pipeline while ensuring local control of each dataset. This open-source distributed pipeline enables biomarker discovery for IO therapies without requiring centralized datasets. We performed univariable analyses to assess the association of 120 GE signatures previously published as TME and IO signatures in pan-cancer, cancer-specific, and treatment-specific contexts. Furthermore, we developed a multivariable predictive model based on IO-associated signatures and evaluated its performance against the pan-cancer PredictIO signature and other locally trained multivariable models, using four independent validation datasets [26]. The distributed pipeline provides a strong foundation for robust and generalizable biomarker discovery and response prediction, demonstrating the effectiveness and interpretability of non-centralized architectures in enhancing biomarker predictive power.

## Materials and Methods

### Immunotherapy datasets

We conducted a systematic review of published clinical studies to identify those that reported both clinical and pre-therapy transcriptomic data for patients treated with IO therapies, including anti-PD-1/L1, anti-CTLA-4, or combination treatments [26]. Transcriptome data were processed using Kallisto 0.46.1 [36] and annotated with Gencode V40 [37]. Log2-transformed TPM (Transcripts Per Million) values were used for protein-coding RNA profiles. When raw FASTQ files were unavailable, raw count or TPM data were downloaded from the publication. For studies providing only FPKM (Fragments Per Kilobase of transcript per Million mapped reads) data, values were converted to TPM [26].

Clinical outcome data included response, as assessed by the Response Evaluation Criteria in Solid Tumors (RECIST) and its derivatives, progression-free survival (PFS), and overall survival (OS). Patients were classified as Responders (R) if they achieved a RECIST (v1.1) complete or partial response, or stable disease (SD) without a PFS event within 6 months. Non-responders (NR) were defined as those with RECIST progressive disease or SD with a PFS event within 6 months. In the absence of RECIST information, patients were classified as R or NR based on PFS events at 6 months. If PFS data were unavailable, patients with RECIST SD as the best response were classified as not assessable. Clinical data were standardized using the mCODE and ICGC-ARGO data dictionaries (**Supplementary File 1**) [38,39].

We collected 21 fully curated, standardized, transparent, and reproducible public MultiAssayExperiment datasets from the cloud-based ORCESTRA platform (orcestra.ca) (**Supplementary Table 1**) [40]. All data were processed in alignment with the FAIR (Findable, Accessible, Interoperable, and Reusable) principles [41]. Each treatment group within a clinical trial and each cancer type were treated as separate units for creating the datasets. For each dataset, genes with zero expression in at least 50% of samples were filtered out. Exploratory analyses were conducted to aid in dataset selection for further analysis, examine feature distribution, and assess heteroskedasticity to identify potential unwanted variations associated with known or unknown batch effects [42] (**Supplementary** Figure 1). All training datasets are publicly available without restrictions, while the four validation datasets have restricted access. These private datasets cover various cancer types and include the INSPIRE (NCT02644369) pan-cancer dataset [43], the Hartwig pan-cancer dataset (NCT02925234) [44], the OAK (NCT02008227) lung cancer dataset [45], the POPLAR (NCT01903993) lung cancer dataset [46], and the IMmotion150 (NCT01984242) kidney cancer dataset [47] (**Supplementary** Figure 1 **and Supplementary Tables 1 and 3**).

### Curation of gene signature and score computation

Published gene expression (GE) signatures, including IO biomarkers and TME-related markers, were used to gain insights into the TME’s cellular composition and functional status to enhance IO therapy [26,48]. These signatures were extracted from the literature and annotated manually using Gencode V40 [37], with HUGO Gene Symbols as the primary identifier linked to Entrez Gene IDs and Ensembl Gene IDs (**Supplementary Table 2**). GE signatures, including unweighted genes, were computed using the Gene Set Variation Analysis (GSVA) enrichment score via the ‘GSVA’ (v.1.50) R package [49]. GE signatures, where genes were assigned specific weights (+1 for upregulated and -1 for downregulated genes), were calculated using the weighted mean expression approach. Scores for signatures requiring specific computational algorithms were obtained from their original publications. Signatures were computed for each study only if at least 80% of their genes were present in the data. After calculating the IO-specific signatures, a z-score transformation was applied to the genes within each signature before applying additional methods.

### Gene signature similarity analysis

To evaluate the similarity between the curated signatures, both common genes and signature scores were analyzed. A matrix of overlapping genes for each signature was computed to assess this similarity. The distance between gene signatures was evaluated using Principal Component Analysis (PCA). Clusters of signatures were identified by applying the first two principal components (PC1 and PC2) to Affinity Propagation Clustering (APC), implemented via the ‘apcluster’ (v.1.4.13) R package. For clusters containing at least two signatures, all genes were combined, and pathway enrichment analysis was performed using the Kyoto Encyclopedia of Genes and Genomes (KEGG) pathways via the ‘enrichR’ R package (v.3.2).

Pearson correlations between the signature scores were computed within each dataset. The correlations were then pooled using the DerSimonian and Laird random-effects meta-analysis with the inverse variance weighting approach, implemented via the ‘meta’ (v.7.0) R package [50,51]. To provide a clear visual representation, hierarchical clustering was applied using the complete linkage method and Manhattan distance. The resulting dendrogram, alongside the Variance Ratio Criterion (VRC or Calinski-Harabasz index) [52], was analyzed to determine the optimal number of clusters. The gene overlap matrix and integrated correlation were visualized using the ‘ComplexHeatmap’ (v.2.18) R package.

### Univariable prediction of Immuno-Oncology response

The association of GE signatures with IO responses was evaluated based on the compendium of public datasets (**Supplementary** Figure 1). Signature associations with IO response (R/NR) were analyzed using a logistic regression model, while associations with IO survival (OS or PFS) were assessed using the Cox proportional hazards model to evaluate survival discrimination. To enhance reproducibility, results from individual independent datasets were pooled using the DerSimonian and Laird random-effects meta-analysis with inverse variance weighting, implemented in the ‘meta’ (v.3.0) R package [50,51]. Heterogeneity across datasets was assessed using the Q statistic and 𝐼 , which quantifies the proportion of total variation due to heterogeneity rather than sampling error [53]. Subgroup meta-analyses were performed, when applicable, to evaluate the impact of clinical variables such as cancer type and type of immunotherapy on IO response prediction. Multiple testing corrections were applied using the Benjamini-Hochberg False Discovery Rate (FDR) method in the ‘stats’ (v.3.6.2) R package [54]. Associations were considered statistically significant for p-values or FDR lower than or equal to 5%. The estimated coefficients, ‘logOR’ and ‘logHR,’ represent the natural logarithms of the odds ratio (OR) and hazard ratio (HR), respectively. OR refers to the odds of non-response, while HR denotes the hazard rate for progression or death.

### Distributed univariable biomarker discovery

We used the cloud-based computational platform Code Ocean [55] to implement a distributed biomarker discovery pipeline, with institutions as nodes and no centralized infrastructure or data (**Figure 1**). This framework involves institutions sharing the same feature space (i.e., gene expression or signatures) but different sample spaces. Datasets from each institution were used, adhering to similar inclusion criteria and formats, including previously described clinical variables and gene expression profiles. At the univariable analysis, the proposed framework was simulated by adapting our existing meta-analysis pipeline to each institution to predict IO response based on curated signatures. The process involves computing the signature score, effect size, variance, and significance of signature associations of interest at each institution, and then transferring these non-confidential data to the global server (or node) to compute the final aggregated or pooled estimates using a meta-analysis approach. The aggregated model results can predict IO response and are designed for interpretability (**Figure 1B**).

**Figure 1:**
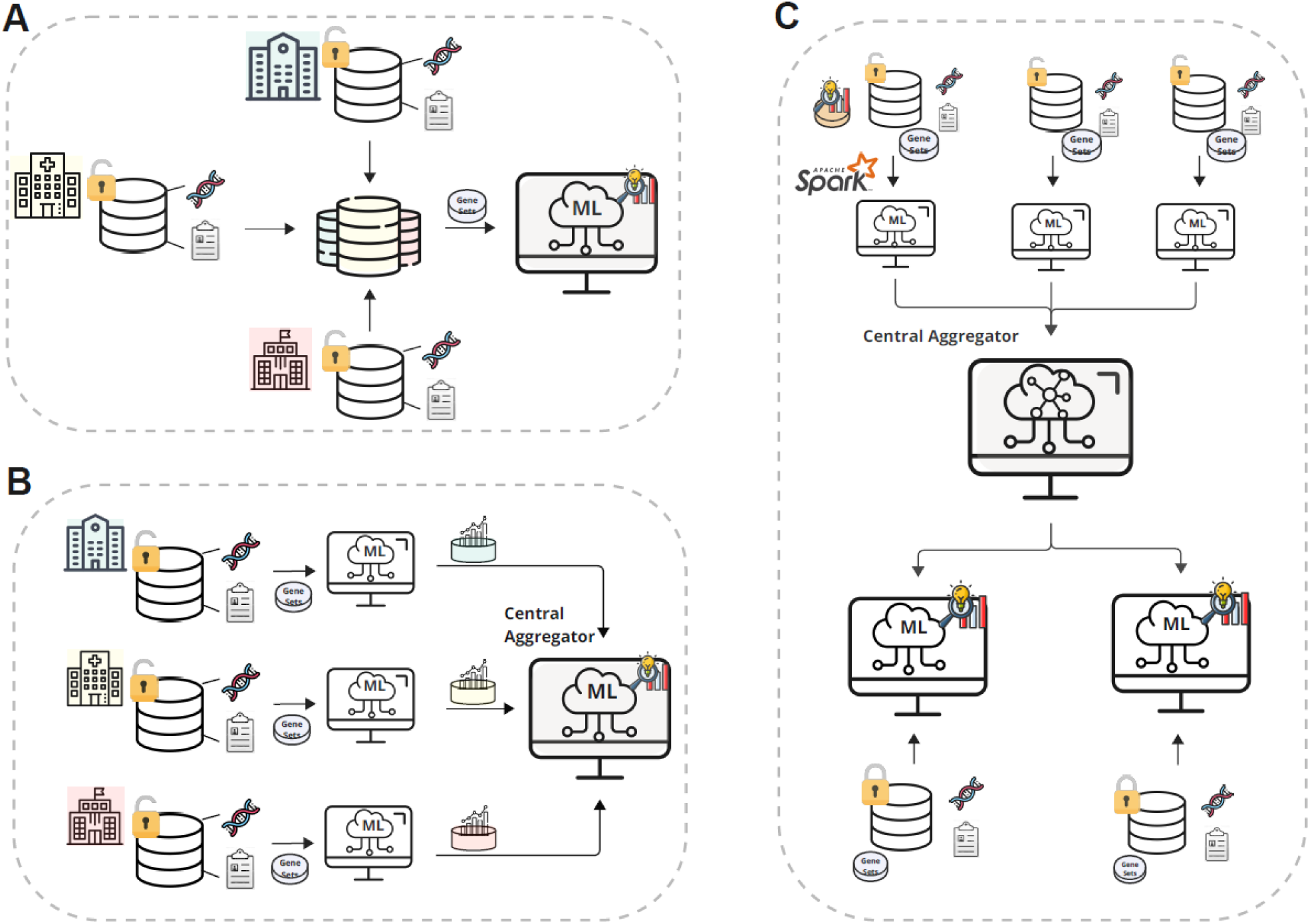
**Cloud-based distributed pipeline for IO biomarker discovery**. (**A**) Centralized analysis where all datasets are transferred to a single location — often impractical due to privacy and policy constraints. (**B**) Distributed association analysis with local computations at each center; only non-sensitive summary data are shared for central aggregation. (**C**) Distributed XGBoost modeling pipeline. Independent models were trained in parallel using Apache Spark, tuned via grid search, and combined into a global ensemble without data pooling. The final model was validated on diverse private cancer datasets.

### Distributed multivariable predictive modelling

The multivariable predictive model was developed using a distributed pipeline implemented across participating institutions and evaluated against two alternative models: the univariable pan-cancer PredictIO signature [26] and multivariable models trained locally on individual datasets. Independent datasets with sufficient patient numbers for modeling were selected and retained within each institution to ensure data privacy (**Supplementary** Figure 1). Distributed model training was performed using Apache Spark and its R API, SparkR (v3.2.1), which enabled parallelized operations across institutional compute environments (**Figure 1C**). At each node, independent XGBoost models were trained using the gradient boosting algorithm (implemented via the ‘xgboost’ R package, v1.7.8.1) without data pooling. To support reproducibility and alignment across institutions, a shared set of IO signatures and standardized training scripts were provided. Hyperparameters were tuned locally using a grid search approach to optimize model performance within each dataset.

A tree-based aggregation strategy was used to combine the independently trained XGBoost models into a single global model at the central node. Following a tree-bagging approach, individual decision trees from each local model were sequentially merged into the global ensemble without altering their learned structures and parameters. This enabled efficient integration while preserving dataset-specific patterns captured during local training. The global model, along with the PredictIO signature and local XGBoost models, was validated using the largest available transcriptomic dataset spanning multiple cancer types (**Supplementary Table 3**). Validation datasets were selected for their diverse composition, including both pan-cancer and single-cancer patients, to ensure robust evaluation of model performance. Comparative analysis was conducted using area under the receiver operating characteristic curve (AUC) with 95% confidence intervals (CIs), computed via the ‘pROC’ R package (v1.18.5).

### Statistical analysis

Descriptive statistics for clinical variables were reported as frequencies and percentages for categorical variables, and as the median and interquartile range (IQR) for continuous variables. Comparisons between categorical variables were assessed using the Chi-squared test or Fisher’s exact test. Differences among more than two groups for continuous variables were assessed using the Kruskal-Wallis test.

### Data availability

The curated clinical and transcriptomic IO data used in this analysis are publicly available at https://www.orcestra.ca/clinical_icb. Access to the private datasets requires submitting a request through the source data repository or its website. The curated IO signatures used in this analysis are publicly available at <u>bhklab/SignatureSets</u> (v1.0).

### Code availability

The simulated univariable and multivariable modelling workflows are available at: PredictioR R package: <u>bhklab/PredictioR</u> (v1.0)

Univariable pipeline: https://github.com/bhklab/PredictIO-UV-Dist Multivariable pipeline: https://github.com/bhklab/PredictIO-MV-Dist

## Results

### Compendium of clinical datasets

We first reviewed the descriptive statistics of clinical characteristics reported across both the discovery and validation datasets. Clinical and gene expression data were compiled

from a total of 1,699 patients with advanced solid tumors, all of whom were primarily treated with immune checkpoint inhibitors (**Supplementary** Figure 2). The main cancer types included lung cancer (34%), melanoma (26%), bladder (14%), and kidney cancer (13%). The majority of patients (91%) received PD-1/PD-L1 therapy, while less than 10% received CTLA-4 therapy alone or in combination with PD-1/PD-L1. Within these datasets, 24% of patients were responders and 57% were non-responders; response data were missing for the remaining 19% of patients. The median PFS and OS were 2.82 months (IQR: 1.45-8.22) and 11.50 months (IQR: 5.13-21.70), respectively (**Table 1** and **Supplementary Table 3**).

**Table 1:** Clinicopathological characteristics of compendium of clinical data. Cancer types named ‘Other’ include Head and Neck (15), Ovary (13), and Breast (11), Prostate (7), Mesothelium (3), and Other (21).

|  |  | All data | Discovery | Validation |
| --- | --- | --- | --- | --- |
|  | N° studies | 23 | 18 | 5 |
|  | N° patients | 1699 | 967 | 732 |
| Age, years | Median (IQR) | 62 (54-69) | 62.5 (55-69) | 55.82 (47 - 65) |
|  | NA (%) | 1353 (80) | 687 (71) | 666 (83) |
| Sex | Female (%) | 414 (24) | 211 (22) | 203 (20) |
|  | Male (%) | 855 (50) | 499 (52) | 356 (35) |
|  | NA (%) | 430 (25) | 257 (26) | 173 (45) |
| Cancer type | Lung (%) | 585 (34) | 144 (15) | 441 (60) |
|  | Melanoma (%) | 447 (26) | 372 (38) | 75 (10) |
|  | Bladder (%) | 233 (14) | 195 (20) | 38 (5) |
|  | Kidney (%) | 223 (13) | 115 (12) | 108 (15) |
|  | Ureteral (%) | 51 (3) | 51 (5) | NA |
|  | Pancreas (%) | 45 (3) | 45 (5) | NA |
|  | Gastric (%) | 45 (3) | 45 (5) | NA |
|  | Other (%) | 70 (4) | NA | 70 (10) |
| Drug treatment | PD-1/PD-L1 (%) | 1547 (91) | 815 (84) | 732 (100) |
|  | CTLA4 (%) | 82 (5) | 82 (8) | NA |
|  | Combo (%) | 70 (4) | 70 (7) | NA |
| Response | No-response (%) | 969 (57) | 517 (53) | 452 (60) |
|  | Response (%) | 412 (24) | 261 (27) | 151 (25) |
|  | NA (%) | 318 (19) | 189 (20) | 129 (15) |
| PFS (months) | No-progression (%) | 278 (16) | 123 (13) | 155 (32) |
|  | Progressed (%) | 760 (45) | 328 (34) | 432 (31) |
|  | NA (%) | 661 (39) | 516 (53) | 145 (37) |
|  | Median PFS | 2.82 | 3.54 | 2.62 |
|  | IQR PFS | 1.45 -8.22 | 2.00-11.17 | 1.38-6.02 |
| OS (months) | Alive (%) | 409 (24) | 270 (28) | 139 (8) |
|  | Dead (%) | 800 (47) | 459 (47) | 341 (27) |
|  | NA (%) | 490 (29) | 238 (25) | 252 (65) |
|  | Median OS | 11.50 | 11.69 | 11.14 |
|  | IQR OS | 5.13-21.70 | 5.34-21.59 | 5.04 - 22.00 |

### Overview of curated RNA signatures

We curated 120 GE signatures previously reported to be associated with the TME and/or sensitivity or resistance to IO therapy, based on publications from 2006 to 2023. Of these, 35 were sensitivity-associated and 18 were linked to resistance. To assess the diversity of these signatures, we applied clustering techniques to identify relationships among the IO and TME signatures separately. For the IO signatures, we identified seven distinct clusters. Four clusters were enriched for pathways associated with chemokine, cytokine, and T-cell receptor signaling, two clusters were related to antigen-presentation machinery, and one cluster was a heterogeneous group enriched for cell cycle processes and pathways in cancer (**Supplementary** Figure 3A). Similarly, for the TME signatures, we observed four clusters. Two clusters, which contained the majority of signatures, were associated with chemokine, cytokine, T-cell receptor signaling, and antigen-presenting machinery. The other two clusters were enriched for cell cycle processes, TGF-beta signaling, p53 signaling, and cancer-related pathways (**Supplementary** Figure 3B). These findings indicate that TME signatures broadly capture the heterogeneity of the tumor microenvironment. The diversity of signature clusters highlights the broad landscape of immune and TME signatures curated in our compendium.

To evaluate relationships and redundancies among signatures, we calculated Pearson correlation coefficients between signature scores, which were derived by summarizing gene expression levels within each signature for each sample. To ensure data privacy, correlations were calculated independently at each institution and then aggregated on a global server. A random-effects meta-analysis was applied to produce pan-cancer estimates, accounting for dataset heterogeneity and providing robust global correlation measures (**Figure 2A**). Signatures for Neutrophils [56], Plasmacytoid Dendritic Cells [57], B Cells, CD56+ Natural Killer Cells, Natural Killer Cells, Effector Memory T Cells, Activated Dendritic Cells, and T Gamma Delta Cells [58] were excluded from the integration step as fewer than 80% of their genes were present in the datasets, below the required threshold for computation. We identified six clusters of signatures: four clusters associated with chemokine, cytokine, and antigen-presenting machinery, and two heterogeneous clusters related to cancer-related pathways. Signatures annotated as IO-sensitive and TME showed strong correlations, while signatures annotated as IO-resistance were weakly correlated (**Figure 2B**).

**Figure 2:**
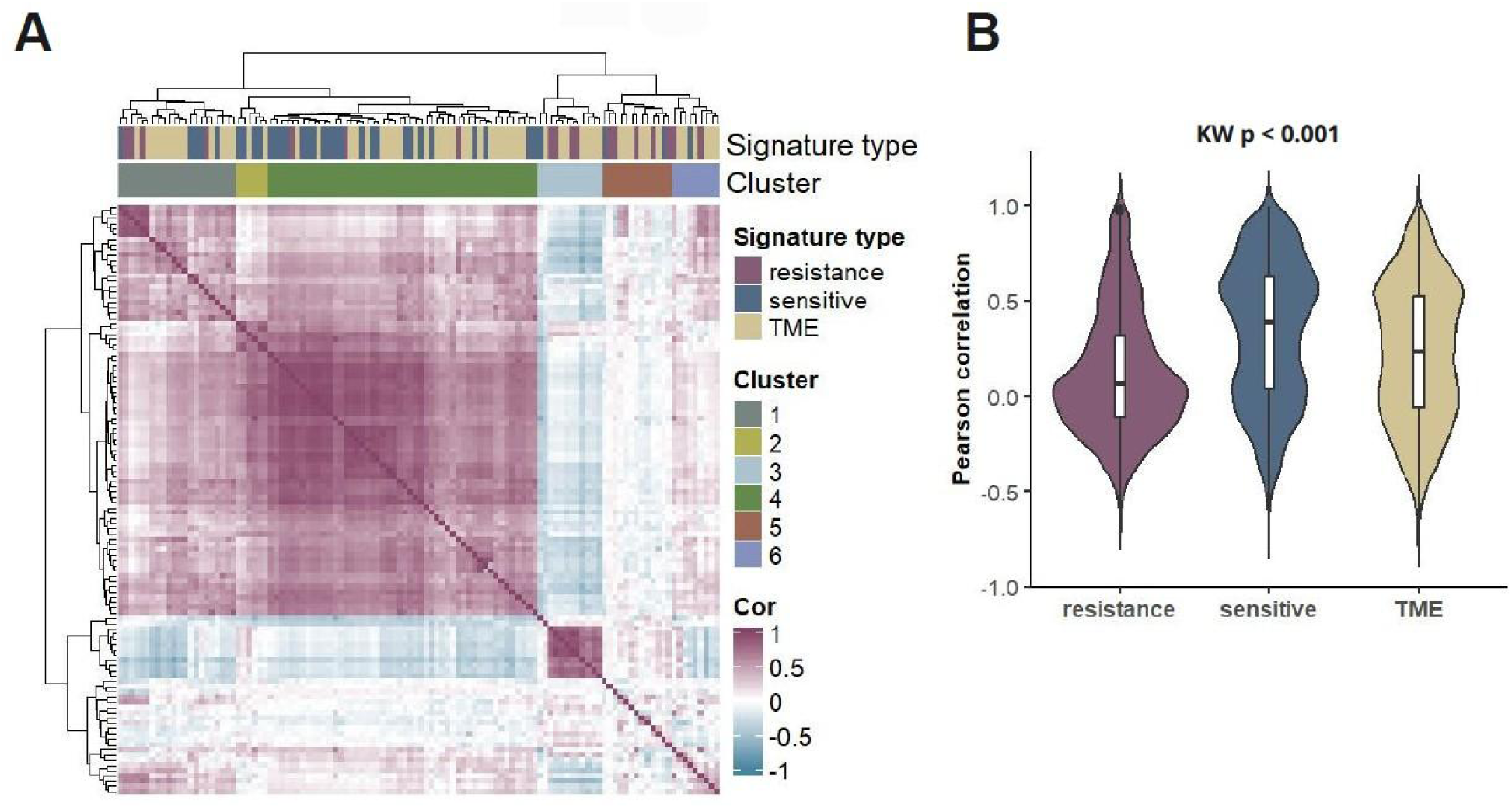
Correlation of the GE curated signatures. (**A**) Heatmap of meta Pearson correlation of the 120 curated GE signatures in the pan-cancer datasets. Hierarchical clustering was performed using the complete linkage method with Manhattan distance. (**B**) Violin plots comparing Pearson correlation levels among resistant, sensitive, and TME signatures in pan-cancer datasets. Differences in correlation levels between signature types were assessed using a two-sided Kruskal-Wallis test.

### Assessment of RNA signatures for IO responses

We examined the association between these signatures and IO responses in a distributed framework by assigning each dataset to a virtual institution. In total, 967 patients across 7 cancer types; melanoma, lung, bladder, kidney, ureteral, pancreas, and gastric; were included in the analysis. To ensure data privacy, association analyses were conducted locally, and only non-sensitive summary statistics were shared with a central node (**Figure 1B**). The aggregated association results were then computed using a random-effect meta-analysis approach, providing robust estimates that accounted for variability across institutions. Out of the 112 signatures, 82 (73%) were significantly associated with at least one IO outcome (i.e., OS, PFS, and response) in the pan-cancer setting after correction for multiple testing (FDR < 5%; **Figure 3** and **Supplementary Tables 5-7**). Among these, 45 signatures were associated with all three IO outcomes (Response, PFS and OS). Of the 35 IO-sensitive signatures, 71% were significantly associated with all clinical outcomes, emphasizing their strong predictive relevance in the context of IO therapy. Out of 67 TME-related signatures, 27% were significantly associated with at least one IO outcome, highlighting their role in modulating immune responses. In contrast, only 2 out of 18 IO-resistance signatures demonstrated significant associations across all IO outcomes, reflecting greater variability in their predictive capacity due to their involvement in diverse biological pathways.

**Figure 3:**
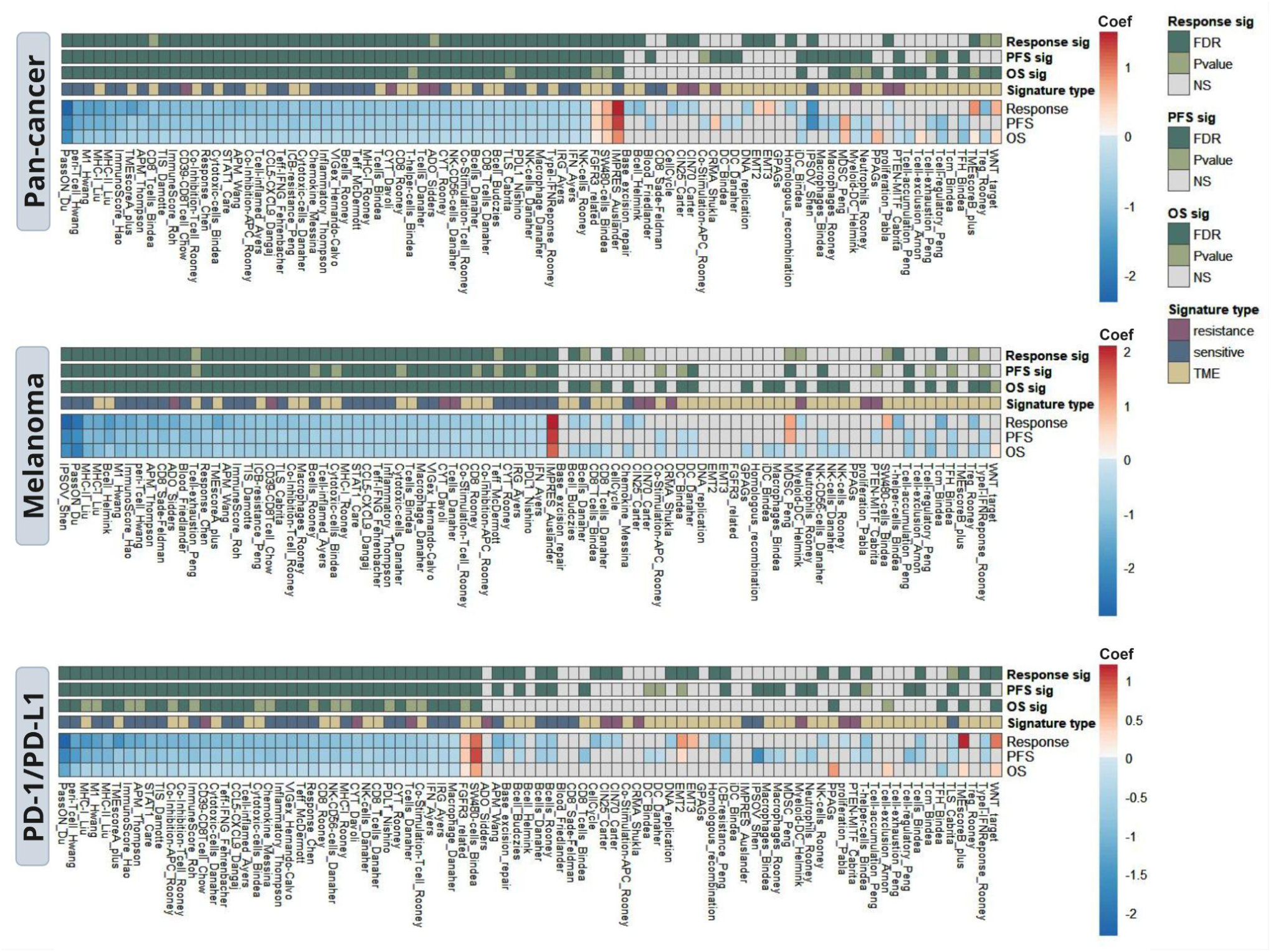
Curated GE signatures are associated with clinical outcomes in IO-treated patients, focusing on those treated with immune checkpoint inhibitors. Meta-federation of GE signatures with IO response, progression-free survival (PFS), and overall survival (OS) in pan-cancer, cancer-specific (melanoma), and treatment-specific (PD-1/PD-L1) settings. The coefficient value in the heatmap corresponds to the pooled logOR or logHR of the meta-analysis. Signatures were identified as “sensitive”, “resistant”, and “TME” based on corresponding publications. Missing coefficient information is shown as grey boxes. P-values were corrected for multiple testing using the Benjamini-Hochberg False Discovery Rate (FDR) correction. Associations with FDR ≤ 0.05 are shown in dark green, while those with an FDR > 0.05 and a P-value ≤ 0.05 are shown in light green. NS: not significant at P-value and/or FDR.

Cancer-specific analyses revealed similar patterns of predictive associations in melanoma compared to the pan-cancer setting, with 30 signatures significantly associated with IO outcomes (FDR ≤ 0.05; **Figure 3** and **Supplementary Tables 8-10**). Notably, 19 IO-sensitive signatures were linked to better OS in melanoma, while the IMPRES signature [59] was the only signature significantly associated with worse OS. In contrast, lung cancer exhibited fewer associated signatures, likely due to the smaller sample size and the reduced number of significant signatures after multiple testing corrections. Stratifying datasets by PD-1/PD-L1 expression revealed a similar pattern to the pan-cancer and melanoma analyses, with 60% of signatures significantly associated with at least one IO outcome (FDR ≤ 5%; **Figure 3** and **Supplementary Tables 11-13**). Of these, 28% of signatures were associated with all three IO outcomes. Four signatures were negatively associated with OS (indicative of improved survival) in the CTLA-4 treatment group but showed no significant association with response or had insufficient PFS data.

### Biomarkers specific to cancer type and treatments

While pan-cancer analysis leverages the large sample size of our data compendium, potential heterogeneity of biomarkers across different cancer types may prevent the identification of cancer-specific biomarkers predictive of immunotherapy response (**Figure 4**). In cancer-specific analyses, we observed similarities in the associated signatures identified across pan-cancer and melanoma analyses for IO outcomes. Notably, certain signatures emerged as significant biomarkers for OS in melanoma and for response in lung and kidney cancers, but were not detected in the pan-cancer analysis (**Supplementary** Figures 4 and 5). Further, there is considerable overlap between findings in pan-treatment analyses and patients treated with PD-1/PD-L1 inhibitors, particularly for PFS and response. However, a subset of biomarkers was uniquely detected in patients treated with CTLA-4 inhibitors and not in the pan-treatment analysis for OS outcomes (**Supplementary** Figures 6 and 7). Melanoma datasets included more patients compared to lung or kidney cancer datasets, and PD-1/PD-L1 treatments were more common than CTLA-4 or combination therapies. These differences in sample size and treatment distribution likely contribute to the observed patterns.

**Figure 4:**
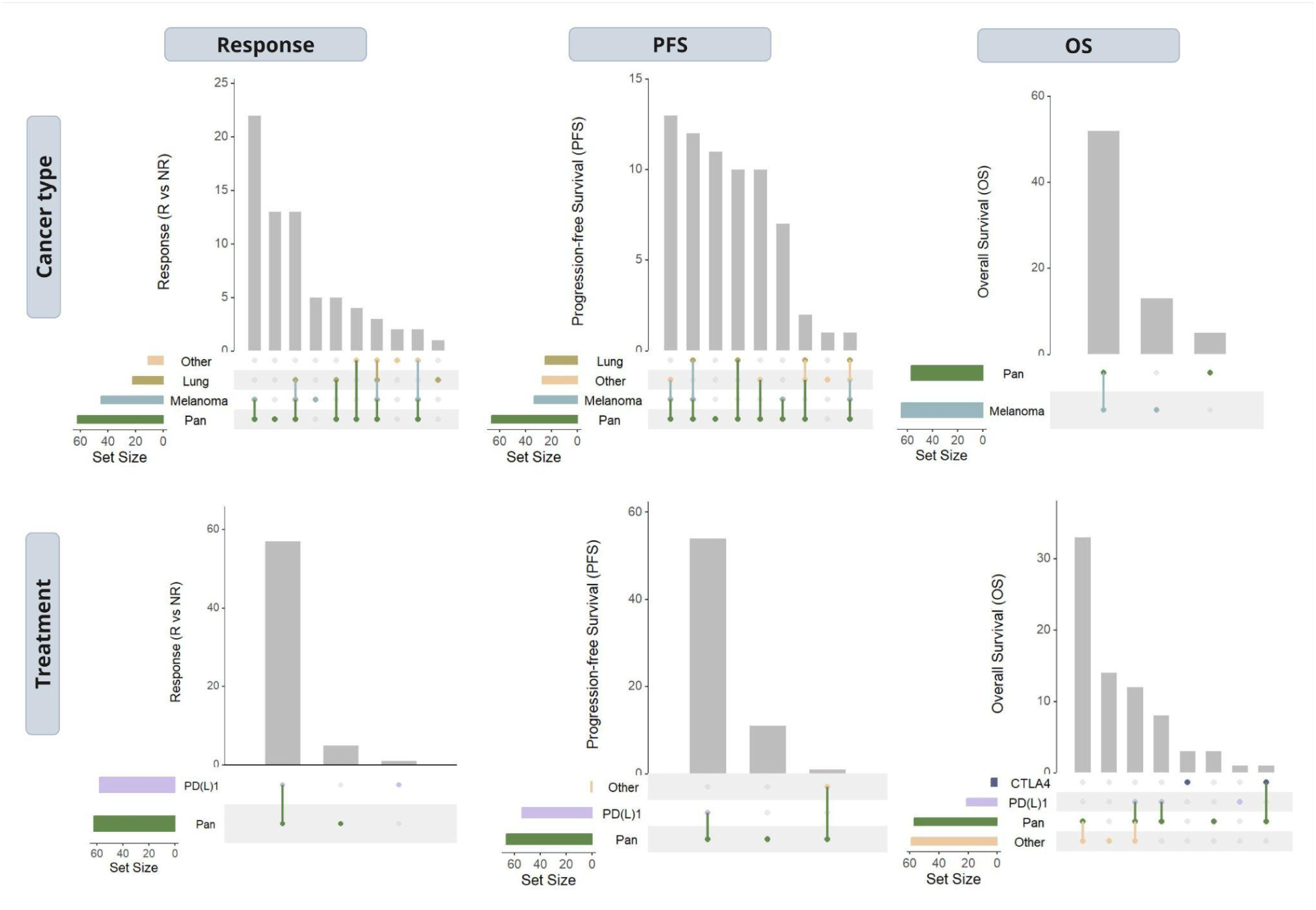
Patterns of associated curated GE signatures with clinical outcomes in specific cancer and treatment. The UpSet plot illustrates the intersections of GE signatures associated with various IO outcomes (response, PFS, and OS) in patients treated with immune checkpoint inhibitors. The analysis includes different cancer types (pan-cancer, melanoma, lung, and kidney) and specific treatments (PD-1/PD-L1 inhibitors and/or CTLA4 inhibitors). Each bar in the plot represents the number of GE signatures shared by the sets indicated by the connected dots below. The size of each bar corresponds to the number of GE signatures common to the connected sets. Only signatures with a False Discovery Rate (FDR) ≤ 0.05 were considered to be significantly associated with IO therapies. The “Other” group represents cancer types or treatments with fewer than three datasets included in the integration analysis.

### A multivariable predictor for IO response

Considering the complexity of the IO biomarker landscape, we investigated whether combining multiple biomarkers into a single non-linear model could improve the prediction of overall IO response likelihood. To achieve this, we trained independent XGBoost models in parallel across 11 datasets comprising 530 patient samples from melanoma, lung, gastric, kidney, and bladder cancers. This distributed framework enabled the development of a multivariable predictive model based on 53 pan-cancer biomarkers commonly present in the datasets (FDR ≤ 0.05; **Figure 3A**). Model performance was compared to two alternatives: the univariable pan-cancer PredictIO signature and locally trained XGBoost models on individual datasets without integration. PredictIO was selected as the benchmark univariable biomarker, as it is a pan-cancer IO signature identified through large-scale gene expression meta-analysis and has demonstrated more consistent predictive performance for IO response than TMB and other published signatures [26].

Following best-practice standards for evaluating diagnostic accuracy, we validated the finalized multivariable predictor using five independent datasets that were excluded from model training (**Supplementary** Figure 1). One of these, the Hartwig dataset, was treated both as pan-cancer and as bladder- and melanoma-specific due to its composition. The PredictIO signature score was computed separately within each validation dataset. For the local models, AUC was averaged across 11 independently trained models, each corresponding to a separate dataset (or node).

The distributed XGBoost model outperformed the average performance of locally trained models across all validation datasets, with AUCs ranging from 0.59 to 0.89 compared to 0.57 to 0.78, respectively. In three of the seven validation datasets, the PredictIO signature slightly outperformed the distributed XGBoost model. These included the pan-cancer INSPIRE dataset (N = 60, AUC = 0.75, 95% CI: [0.61–0.75]), the lung cancer dataset POPLAR (N = 70, AUC = 0.69, 95% CI: [0.51–0.69]), and the kidney cancer dataset IMmotion150 (N = 54, AUC = 0.60, 95% CI: [0.42–0.60]) (**Figure 5**). In the remaining datasets, the distributed XGBoost model outperformed both the PredictIO signature and locally trained models. Overall, these findings demonstrate that the distributed XGBoost model consistently outperformed local models on average, suggesting improved generalizability through distributed training. They also underscore the variability in predictive performance across cancer types and dataset composition.

**Figure 5:**
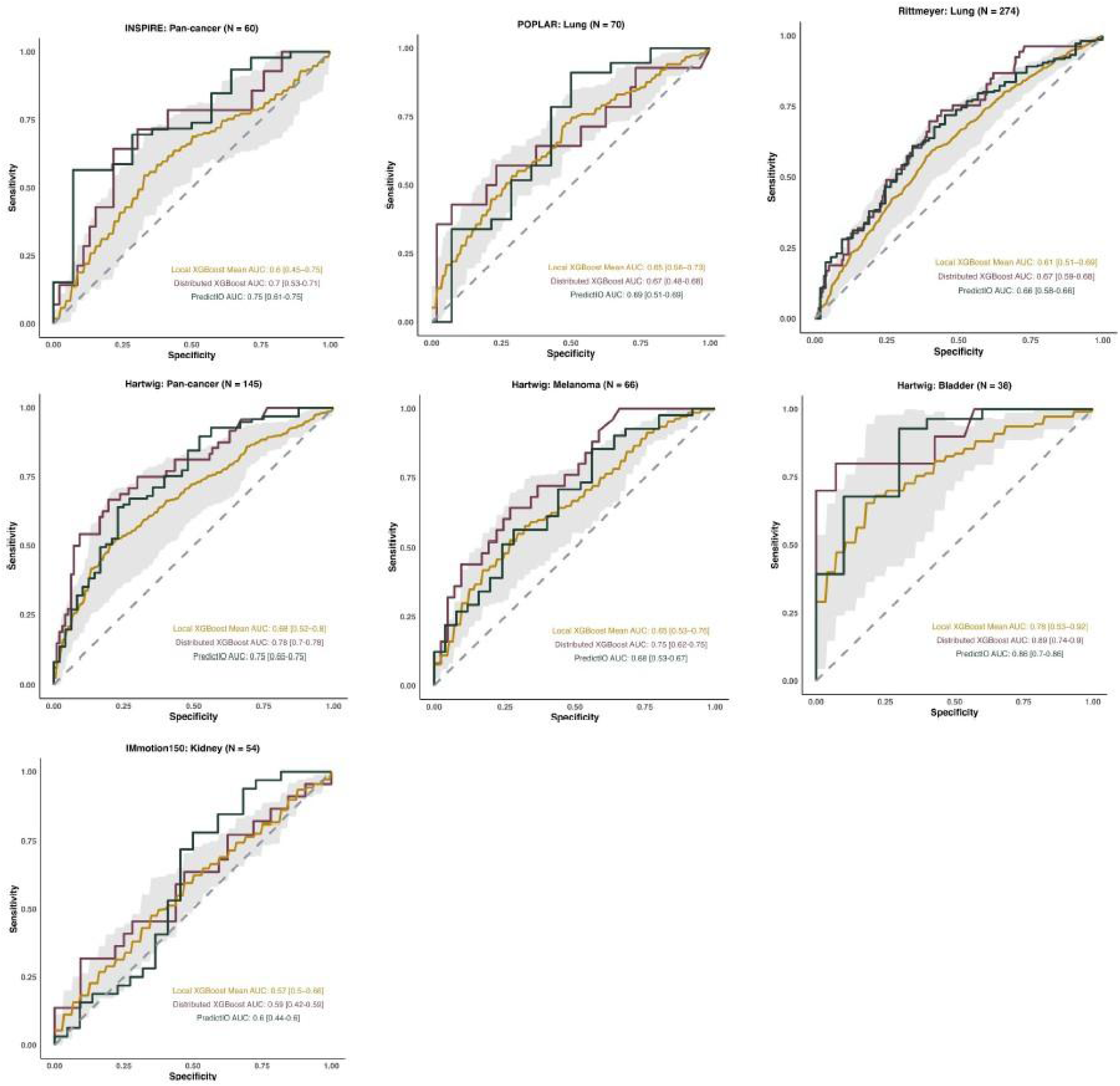
Comparison of distributed, local, and PredictIO-based predictive models. Validation includes ROC curves and AUC values with 95% confidence intervals across four independent datasets (excluded from training). The multivariable distributed model was benchmarked against both the univariable PredictIO signature and multivariable models trained locally on individual datasets, with AUC averaged across the 11 local models.

## Discussion

This study introduces a distributed pipeline for identifying predictive IO biomarkers, with a focus on immune checkpoint inhibitors, while addressing critical data privacy challenges. Unlike traditional centralized approaches, this framework enables seamless multi-institutional collaboration without sharing sensitive data, overcoming key barriers to data integration in biomedical research. By employing this distributed approach, we demonstrated the feasibility of robust biomarker discovery across diverse clinical datasets, ensuring both data privacy and security.

Our distributed meta-integration analysis highlights the importance of cancer- and treatment-specific approaches, with unique biomarkers identified for melanoma, lung cancer, and CTLA-4 therapies, underscoring the need for precision strategies tailored to clinical context. To address the complexity of IO biomarker prediction, we also developed a multivariable model using a distributed training pipeline. In validation, this model outperformed local models on average, suggesting improved generalizability. However, PredictIO demonstrated stronger predictive accuracy in most datasets, consistent with its design as a robust pan-cancer biomarker. These findings highlight both the potential and variability of distributed modeling, influenced by tumor type and dataset composition.

While our distributed pipeline enhances privacy and enables collaboration across institutions, it also presents several important limitations. A key challenge is managing data heterogeneity, including variations in sample processing, sequencing platforms, and clinical variables, which can affect model robustness and comparability across sites [60]. In addition, the approach also places a computational burden on local infrastructure, which may limit scalability in resource-constrained environments. While the framework avoids direct data sharing, recent studies suggest that even aggregated outputs could potentially leak sensitive information under certain attack models, underscoring the need for enhanced privacy safeguards such as secure aggregation or differential privacy [61]. Lastly, the absence of centralized data access can introduce technical challenges that affect training efficiency and complicate the development process as the lack of direct access to data render troubleshooting challenging [60].

Future work should leverage federated pipelines to access a broader range of molecular and clinical data across institutions, including genomics, imaging, spatial profiling, and histopathology. Integrating these complementary modalities can enhance biological resolution and support the discovery of more robust, context-specific biomarkers [62–64]. Incorporating key clinical variables such as age, sex, histology, and cancer stage, will further improve model accuracy and clinical relevance. For broader adoption, scaling the pipeline to larger, real-world datasets and extending it to multi-omics integration will be essential. As immunotherapy increasingly involves the combination of strategies, future efforts must also address the added computational complexity of biomarker development in this evolving landscape.

In conclusion, our study presents a distributed informatics framework for IO biomarker discovery, enabling the integration of clinical and molecular data while safeguarding patient privacy. This distributed pipeline enables federated analysis across institutions, supporting the development of robust, context-specific predictors for different cancers and therapies. By advancing biomarker-driven patient stratification, this approach holds strong potential to improve response rates and ultimately outcomes in precision immuno-oncology.

## Lead contact

Further information and requests for resources and reagents should be directed to and will be fulfilled by the Lead Contact, Benjamin Haibe-Kains.

## Authors’ Contributions

F.A.A. contributed to writing–original draft, writing–review & editing, supervision, investigation, project administration, methodology, formal analysis, validation, software, visualization, data curation, and conceptualization. K.N. contributed to writing–review & editing, formal analysis, validation, software, and visualization. M.N. contributed to writing–review & editing and data curation. K.X.W. contributed to writing–review & editing, investigation. S.K.N. contributed to writing–review & editing, data curation, and conceptualization. N.B. contributed to writing–review & editing, data curation and visualization. N.S., S.S., J.D., N.R., T.L., A.L., R.Z.M., S.N., A.V., R.A., J.S., M.F., and C.M. all contributed to writing–review & editing and conceptualization. B.H.K. contributed to writing–review & editing, supervision, investigation, project administration, methodology, formal analysis, validation, software, visualization, and conceptualization.

## Supporting information

https://docs.google.com/spreadsheets/d/1qN7ix5pjFQA4I9lDW2TTGl_pM8_y4MQH/edit?usp=sharing&ouid=102641196667880928861&rtpof=true&sd=true

## Acknowledgments

We are grateful for the collaborative support provided through the imCORE Network. We thank Iain Buchanan and Simon Adar from Code Ocean for their support setting up the distributed Code Ocean capsules to ensure full transparency and reproducibility, while maximizing the reusability of the biomarker discovery pipelines.

## Supplementary Figures

**Supplementary Figure 1:**
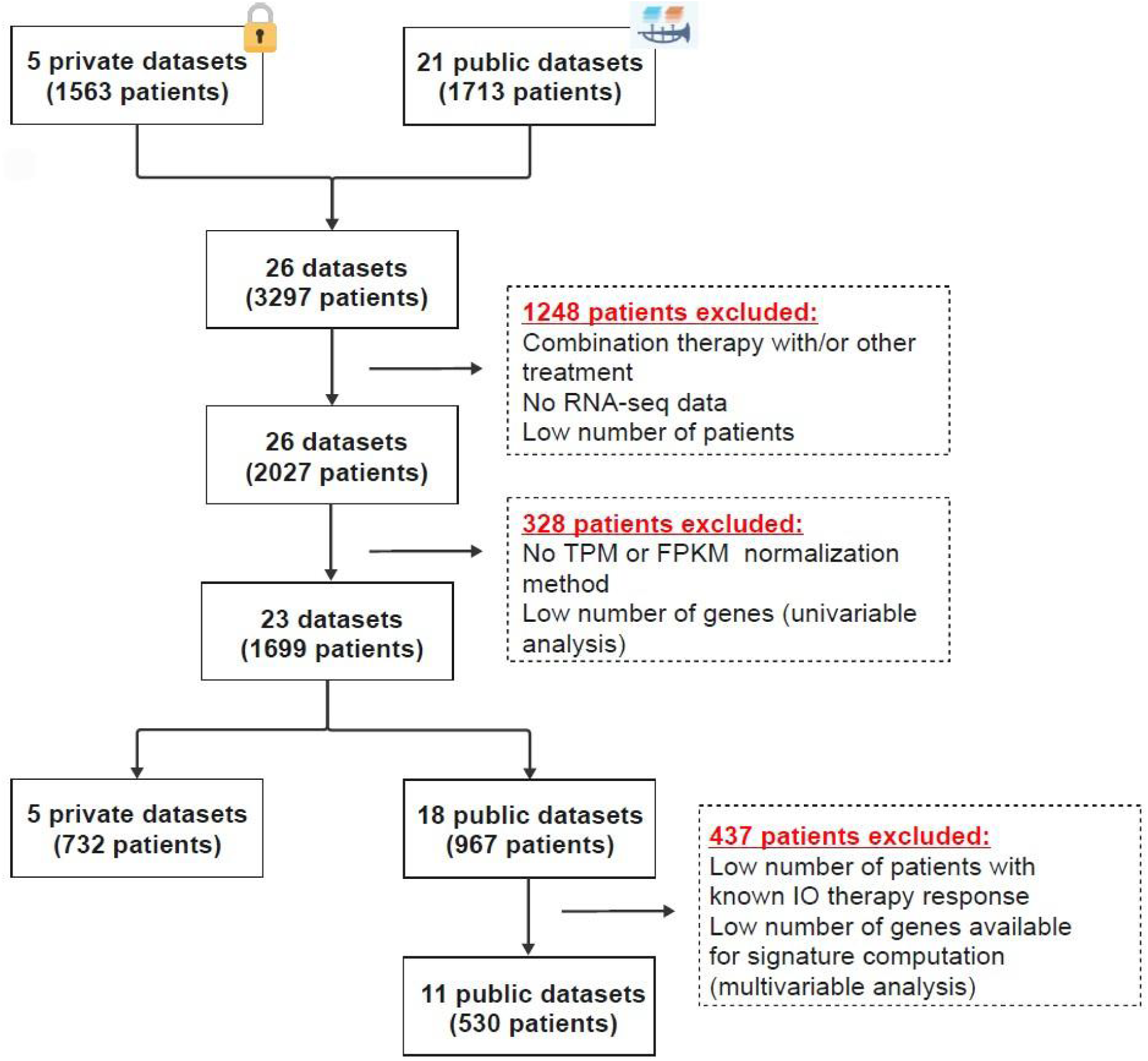
Flowchart of public and private IO datasets used for biomarker discovery.

**Supplementary Figure 2:**
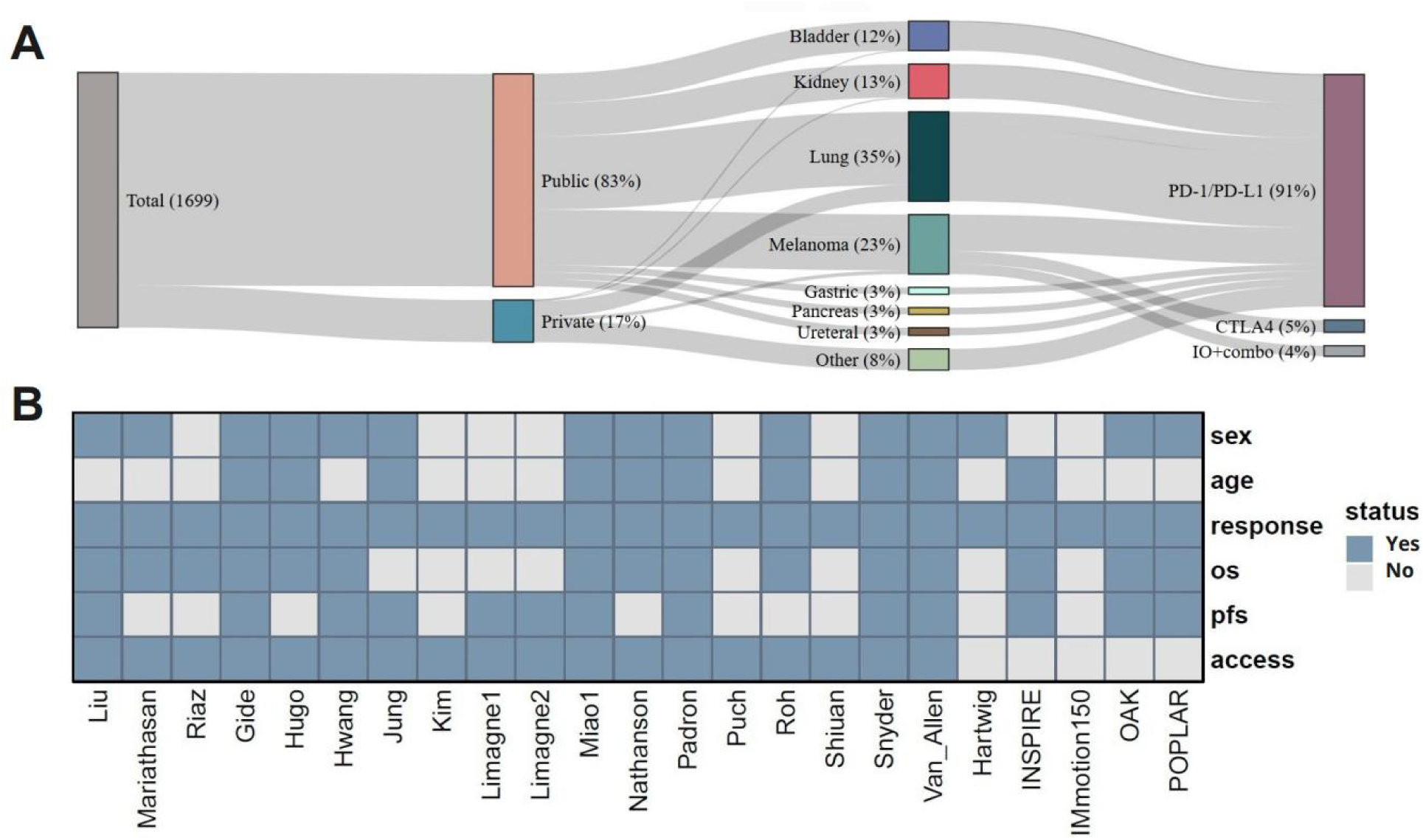
Flow diagram of clinicopathological characteristics and clinical data distribution. (**A**) Sankey diagram illustrating the flow and distribution of clinical data across key variables. (**B**) Heatmap representing the distribution and completeness of clinical data across key variables, highlighting patterns in data availability and variability across the cohort.

**Supplementary Figure 3:**
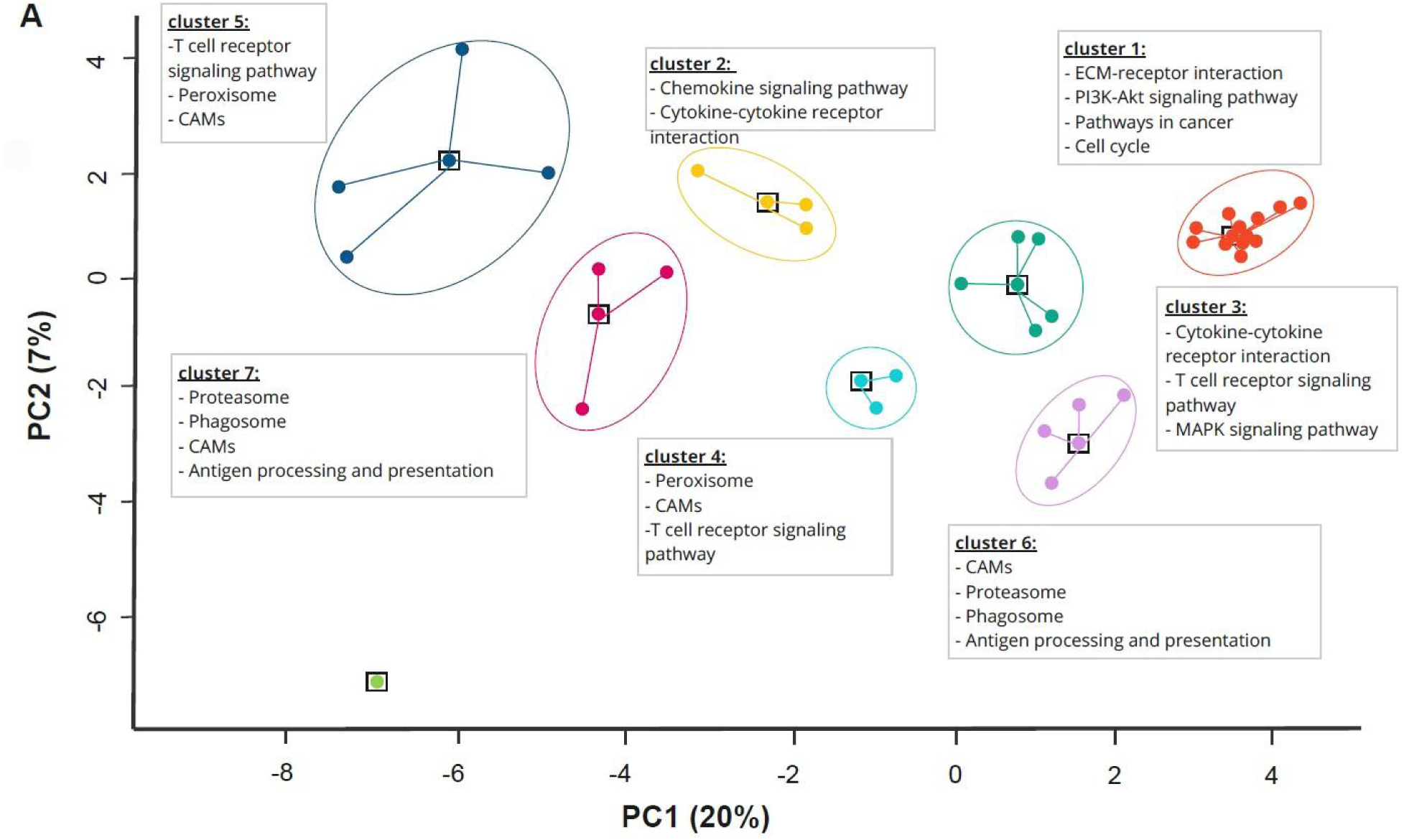
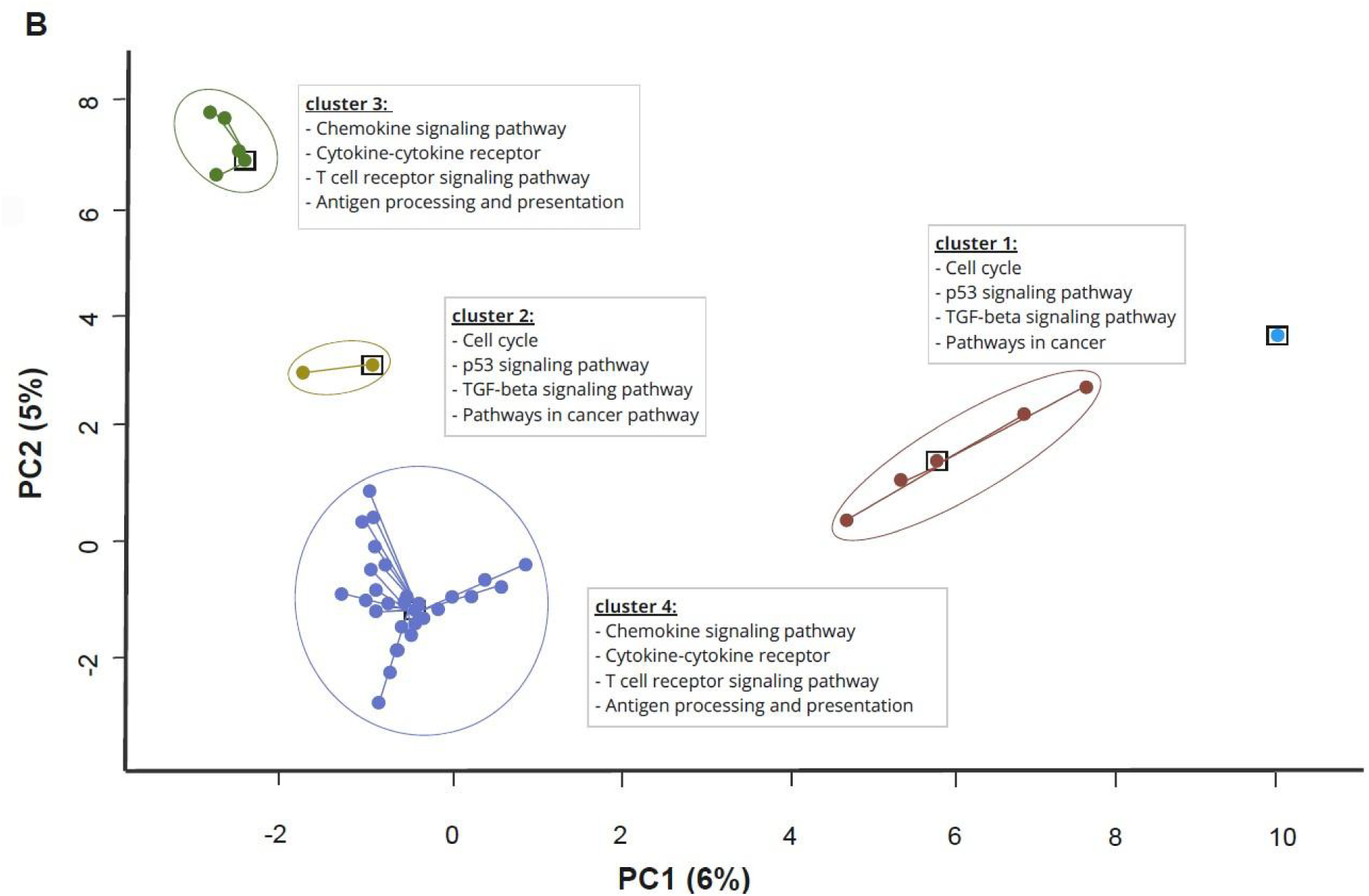
Pattern of the GE curated signatures. Principal component analysis (PCA) of the overlapping genes between each curated (**A**) IO and (**B**) TME signature. Clusters were determined using Affinity Propagation Clustering. The Kyoto Encyclopedia of Genes and Genomes pathway enriched within each cluster was computed from enrichR.

**Supplementary Figure 4:**
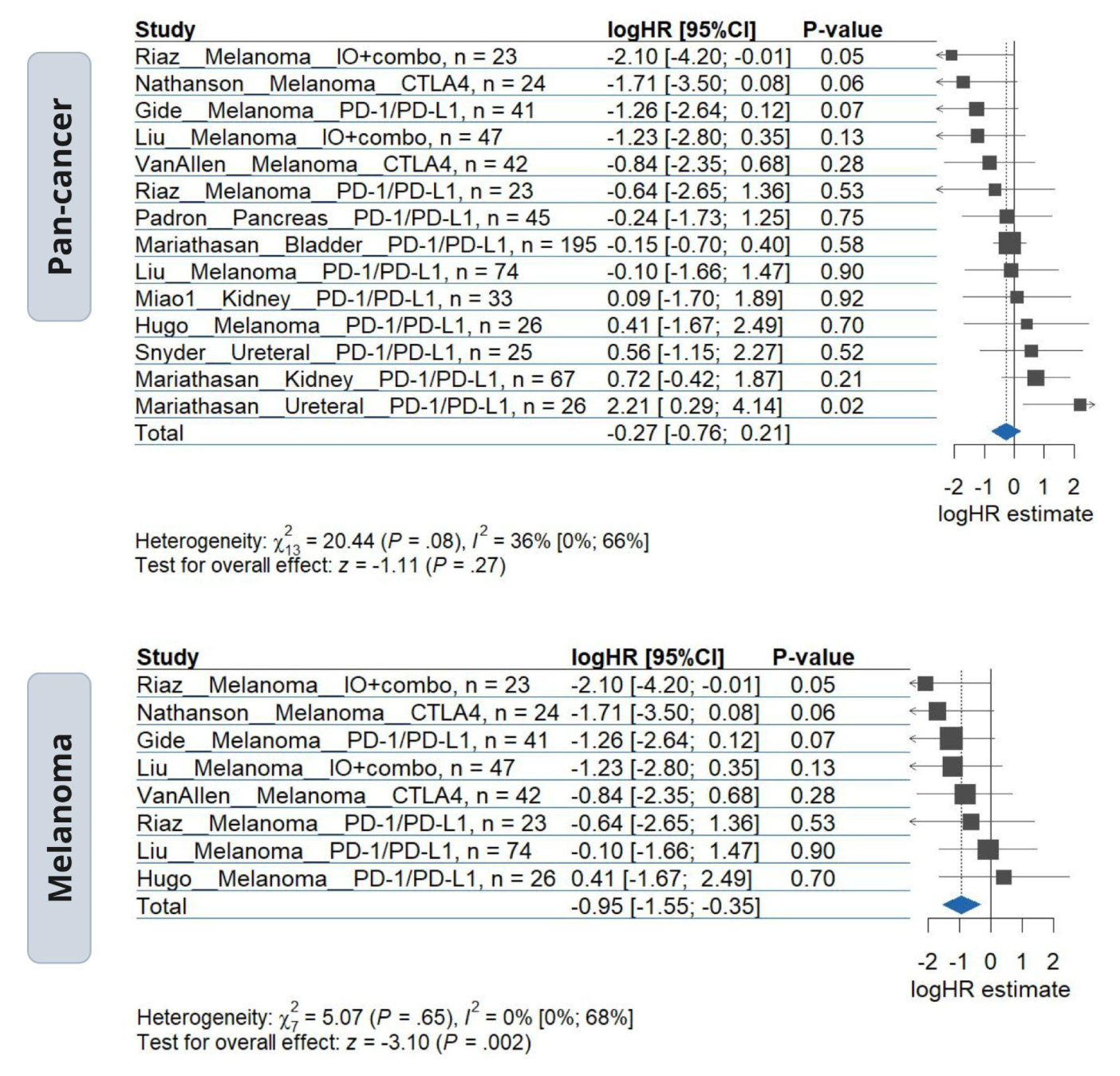
T Cell-exhaustion signature association with overall survival in IO-treated patients. Meta-analysis of T Cell-exhaustion with IO overall survival (OS) as pan-cancer and melanoma analyses. Forest plot displaying the log hazard ratios (logHR) and 95% confidence intervals (CI) for the variable association with OS outcome. Horizontal bars represent the 95% confidence intervals of effect size. The blue diamond represents the overall effect of the variable in IO-treated patients.

**Supplementary Figure 5:**
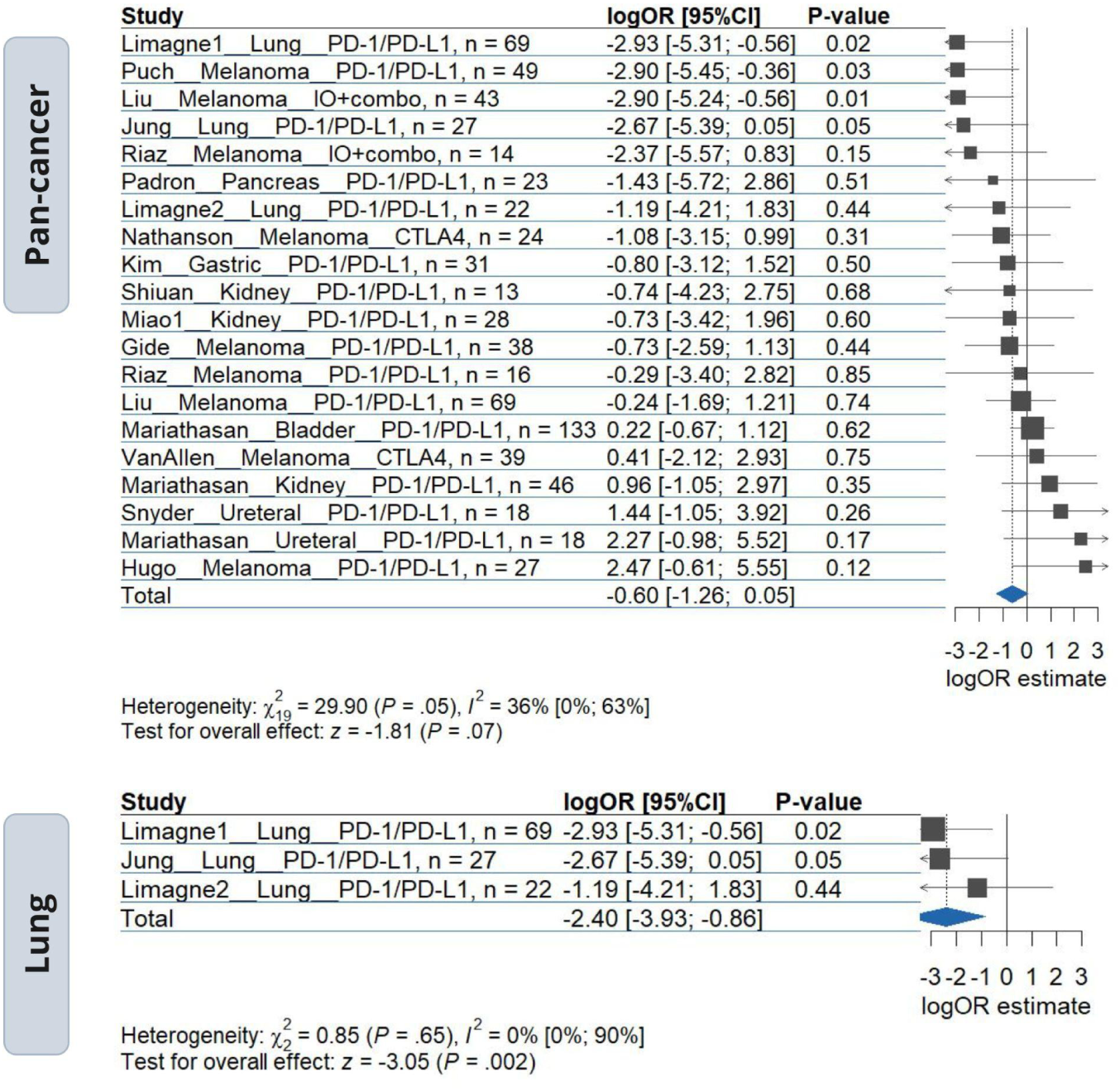
Macrophages_Bindea association with response in IO-treated patients. Meta-analysis of Macrophages_Bindea with IO response as pan-cancer and lung analyses. Forest plot displaying the log odds ratios (logOR) and 95% confidence intervals (CI) for the variable association with response outcome. Horizontal bars represent the 95% confidence intervals of effect-size. The blue diamond represents the overall effect of the variable in IO-treated patients.

**Supplementary Figure 6:**
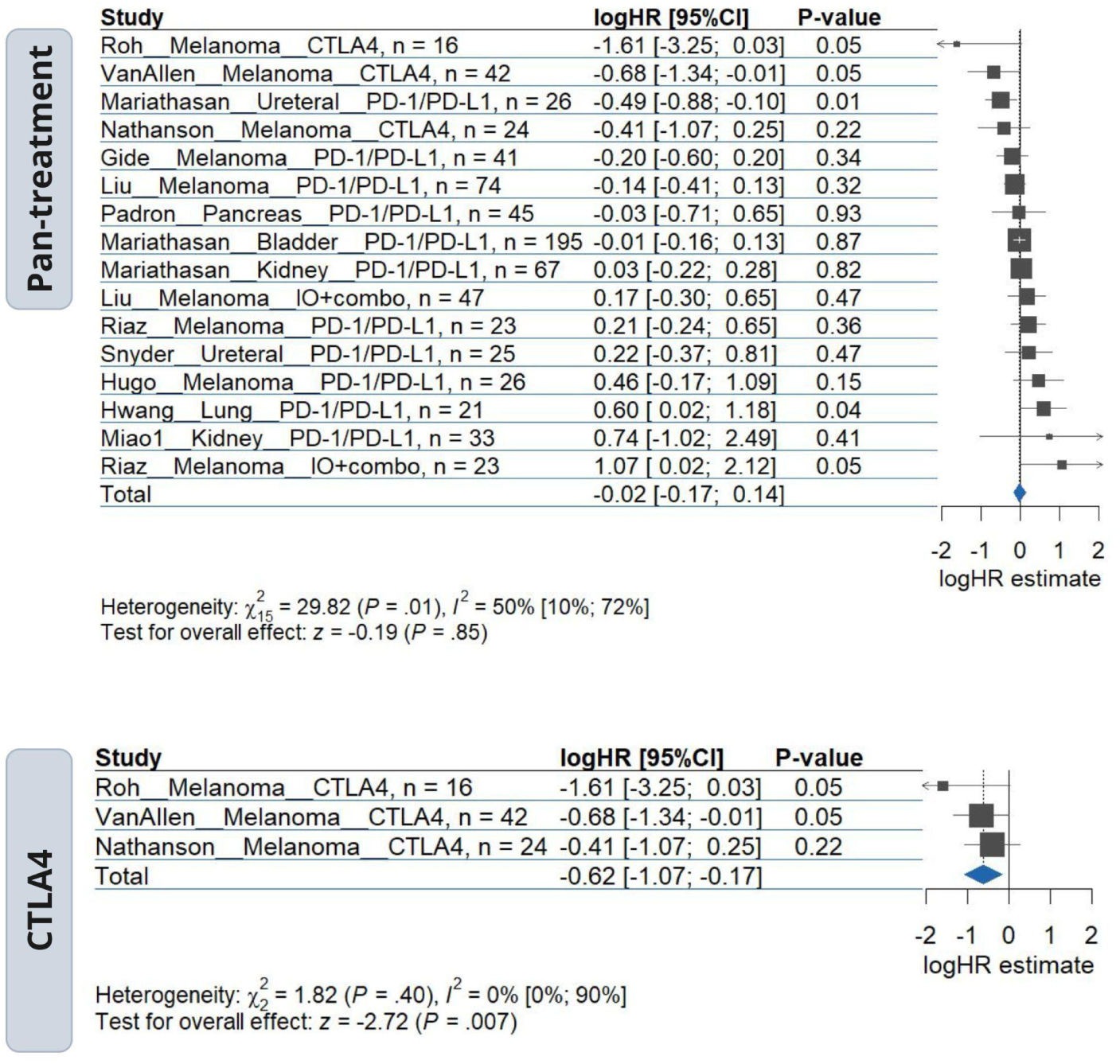
COX-IS_Bonavita association with overall survival in IO-treated patients. Meta-analysis of COX-IS_Bonavita with IO overall survival (OS) as pan-cancer (or pan-treatment) and CTLA-4 analyses. Forest plot displaying the log hazard ratios (logHR) and 95% confidence intervals (CI) for the variable association with response outcome. Horizontal bars represent the 95% confidence intervals of effect-size. The blue diamond represents the overall effect of the variable in IO-treated patients.

**Supplementary Figure 7:**
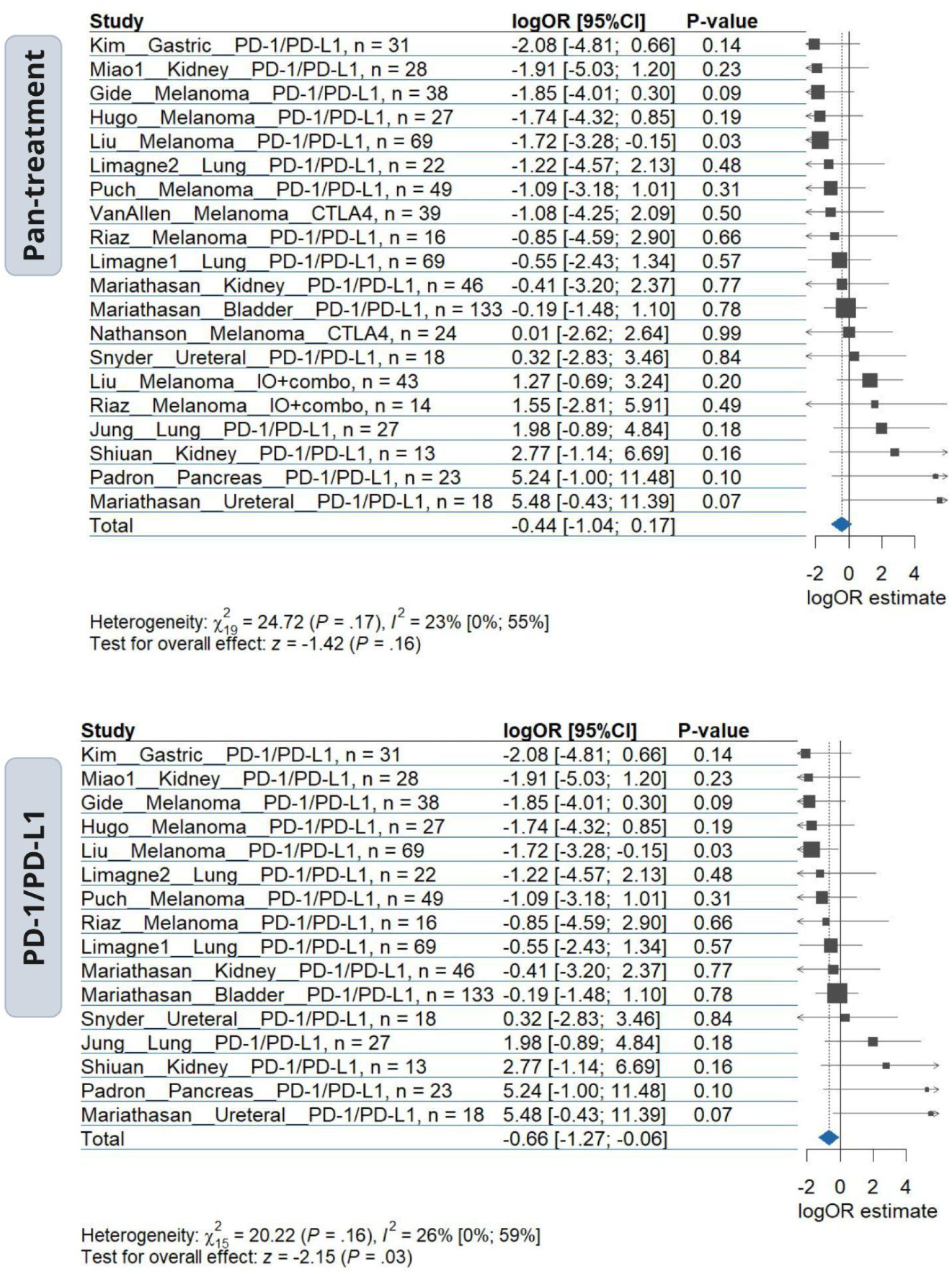
Cell Cycle regulatory signature association with response in IO-treated patients. Meta-analysis of Cell Cycle regulatory signature with IO response as pan-cancer (or pan-treatment) and PD-1/PD-L1 analyses. Forest plot displaying the log odds ratios (logOR) and 95% confidence intervals (CI) for the variable association with response outcome. Horizontal bars represent the 95% confidence intervals of effect-size. The blue diamond represents the overall effect of the variable in IO-treated patients. CellCycle_Reg

